# Electrophysiological measures of sleep pressure during wakefulness in the course of isolation at the Concordia Antarctica station and physical activity as a countermeasure

**DOI:** 10.1101/516567

**Authors:** Gaetan Petit, Vera Abeln, Leopold Summerer, Stefan Schneider, Reto Huber

## Abstract

Concordia station in Antarctica is one of the most remote human outpost on Earth. Because of its geographical location, a winterover at Concordia shares a lot of stressors with a space mission. Following a recent study on the markers of sleep pressure during wakefulness on board of the International Space Station, we were investigating if long term isolation in a space analogue station shows similar effects on sleep pressure. Unlike in space, markers of sleep pressure did not increase during isolation’s constant darkness period in Antarctica. When measures of sleep pressure were high in the evening, psychological strain was increased, emphasising the importance of keeping sleep pressure under physiological levels during the mission. As a first indication for a countermeasure, we showed that one hour of bicycle exercise during lunch time could decrease sleep pressure. All these observations need to be further studied in a more controlled environment.

## Introduction

Sleep quality and sleep quantity are regulated by two interacting process, the homeostatic and the circadian process (1): i) The homeostatic process builds up with time awake. Good sleep quality is defined by an efficient dissipation of sleep pressure overnight which in turn decreases the propensity to fall asleep and improve alertness the following day (2; 3). ii) The circadian process is oscillating on a 24 hours cycle (4). The master clock is regulated by the suprachiasmatic nucleus and constantly adjusted by environmental cues (5). Daylight is one of the main cues to the master clock. Circadian rhythms exert a strong impact on sleep quality and sleep timing (6). Interestingly, EEG theta activity (5-7Hz) in the waking EEG (1) reflects both the homeostatic and the circadian process. The circadian impact can be seen by diurnal fluctuations of theta activity, peaking in the afternoon and again at bed time (7). The homeostatic component, on the other hand, becomes obvious after prolonged wakefulness (i.e. sleep deprivation), which is associated with a significant increase in theta power. (8; 9; 10; 11; 12).

Astronauts are subject to very irregular schedules and workloads (e.g. launchings or dockings of a vehicule) and while orbiting Earth, the International Space Station (ISS) crew witness a sun rise every 90 minutes which can lead to a misalignment of their 24-hour sleep-wake cycle (13). Accordingly, chronic sleep restriction has been reported during space missions (14). We recently provided first evidence for increased of sleep pressure markers during wakefulness in space, i.e measured waking EEG in the theta frequency range, and showed that reaction times were slower when markers of sleep pressure were high (15). To understand the cause of changes in markers of sleep pressure in space we need to disentangle the effect of each space environment stressor on the human brain.

The Concordia research station shares many stressors with long-term space missions and therefore serves as a useful analogue habitat for research on human physiology and psychology (Fig. 1.a). In addition to the harsh environment, Antarctica Concordia station is one of the most remote habitat on Earth, with extreme geographical and social isolation. When temperature drops during the winter (Fig. 1.b), air planes cannot reach the station and all participants are in total isolation for eight months. This total isolation particularity adds an additional psychological stress, since the expedition cannot be stopped in case of failure. Moreover, the 80 scientists from the summer crew are reduced to less than 15 during the winterover. Because of its geographical situation, Concordia station crews do not have any daylight for four months during the winter (May-August). Furthermore, the station is at high altitude (3200m) which implies that the crewmembers have to cope with lower oxygen levels (hypoxia) and low air pressure (hypobaria). Because of its location at the South Pole, the atmospheric pressure is equivalent to an altitude of 3800m outside the polar circle (16). Since the oxygen concentration is at 13% instead of the usual 21%, the blood saturation in oxygen is lower, which increase the heart rate. Hypoxia has shown to impact cognitive performances and sleep quality (17; 18; 19; 20). Fortunately, the adaptation process to high altitude takes place within a few weeks and participants should be acclimated to their new environment for their first measurement (16; 21). Nevertheless, high altitude at Concordia station is a good model of hypoxia and hypobaria conditions encountered during space missions (22).

**Figure 1:**
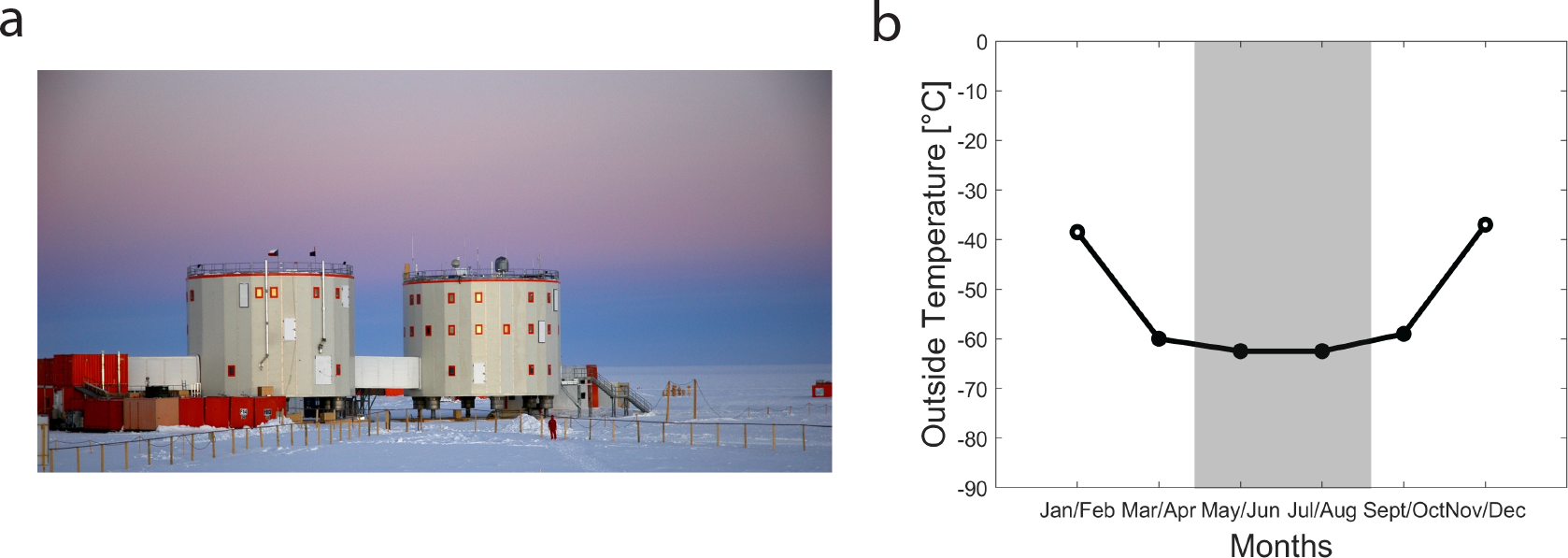
Eight months of isolation at the Concordia Antarctica station. (a) Concordia Antarctica station can host 80 researchers during summer time and usually hosts a crew of 15 researchers over the winter. (b) Mean temperature between 2005 and 2015 at the station. Temperature drops during the isolation period. The black dots represent the four measurements used in this study (Mar/Apr, May/Jun, Jul/Aug and Sept/Oct). The gray shadow marks the four months of constant darkness at the station during the winter period, also called the “permanent midnight of winter”. (credit: ESA).

A complete review of sleep investigations in Antarctica has been recently published (23). Overall, a sleep quality decrease and a delay in the circadian phase has been reported in Antarctica (23). However, a high interindividual variability in the acclimatisation to Antarctic environment was reported in all studies. Interestingly, an actigraphy study from both, the coastal Dumont d’Urville station and Concordia station (3200m), showed that sleep disturbances are more pronounced at high altitude and aggravating during the winter (24).

A first aim of this study is to assess long term isolation’s effect on measures of sleep pressure using wake EEG data recorded at the Concordia Antarctica research station. The benefit of a good night of sleep is well known and chronic sleep restriction comes with a high cost (25; 26; 27; 28; 29). Prolonged wakefulness and an increase of theta activity has been associated with an increase of subjective sleepiness, a decrease of alertness and a degradation of cognitive performances (11; 12; 30; 31).

In Antarctica, physical exercise has shown to improve psychological strain in long term isolation (32). Beside enhancing fitness, strength and compensating for the lack of gravity in space (33), physical exercise countermeasures could also be implemented for cognitive and psychological reasons (34). Interestingly, a recent study in mice showed that wheel running prolonged the waking bouts of the animals without increasing sleep pressure in the motor and somatosensory areas (35). Thus, a second aim of this study is to analyse the relation of sleep pressure markers during wakefulness (i.e. theta activity) with subjective psychological strain and to explore physical exercise effects as a potential countermeasure to dissipate sleep pressure during wakefulness.

## Results

### Theta power across the day

Because of its circadian and homeostatic component, theta power (5-7Hz) oscillates during the day. Thus, in a first step we tested time of day differences in our recordings. Therefore, we defined two groups; a Noon session with five participants (starting between 10h and 13h) and an Evening session with seven participants (starting between 15h and 20h). When comparing the Noon and the Evening recordings, at electrode C3, we observed an increase in the power spectrum spanning lower frequencies from 2.6 to 7.4 Hz (two sample t-test for each (0.125Hz frequency bin), black dots for uncorrected p-values<0.05, n=14 recordings at Noon and n=18 recordings in the Evening) (Fig. 2.a). We then compared only Theta power at the same electrode. Theta power appears higher at Noon than in the Evening (two sample t-test, t=4.693, p<0.001, df= 30, n=14 recordings at Noon and n=18 recordings in the Evening) (Fig. 2.b).

**Figure 2:**
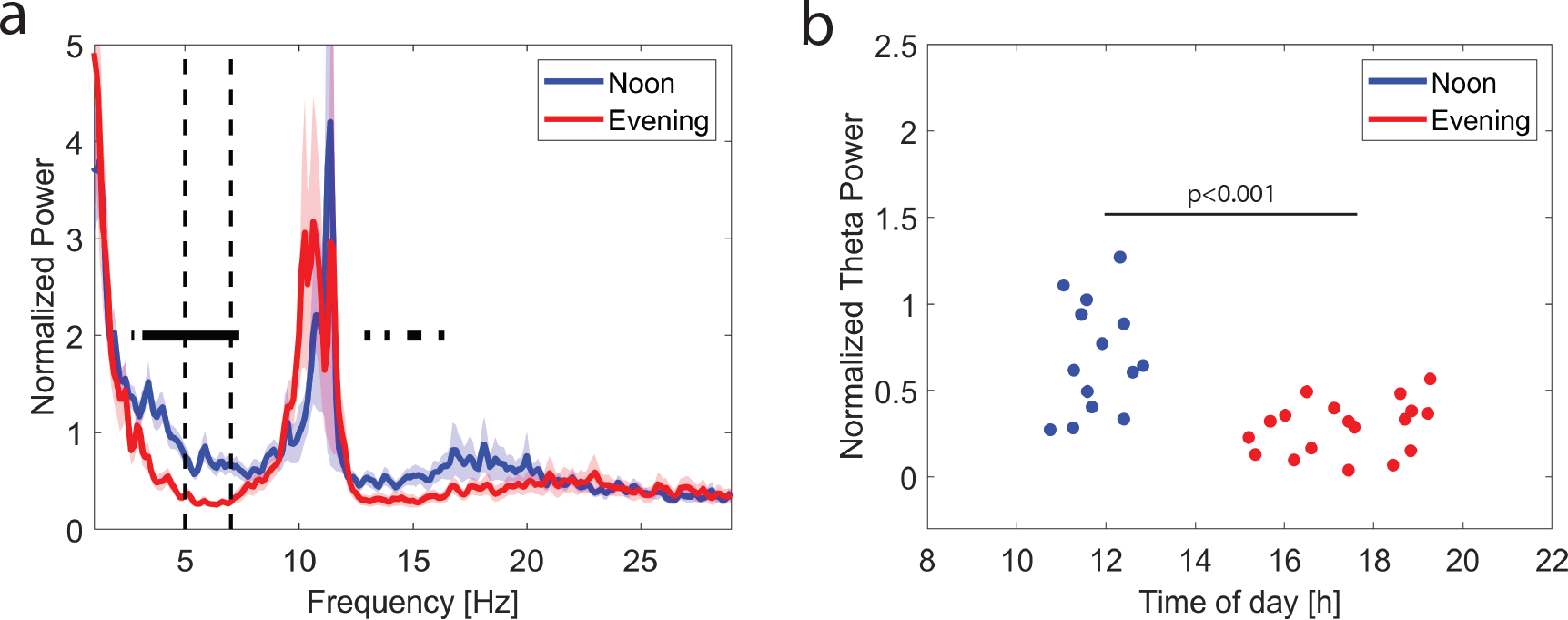
Noon and Evening recordings. (a) The power spectrum at electrode C3 for Noon and Evening recordings (*mean* ± *sem*). Significant decrease of activity from Noon to Evening in the (2.625-7.375Hz) range (black dots for the uncorrected p-values<0.05). (b) The blue dots represent theta power at electrode C3 for recordings taken at Noon (n=14 recordings for 5 participants) and the red dots represent recordings taken in the Evening (n=18 recordings for 7 participants). At electrode C3 theta power is higher at Noon than in the evening.

### Theta power during isolation period

Each participant was recorded every six weeks during the isolation winter, defining four isolation conditions: Mar/Apr, May/Jun, Jul/Aug and Sept/Oct. We represented the topographical distributions of theta power over the scalp for each isolation condition at Noon and in the Evening (Fig. 3.a). Theta power was most pronounced over central and the occipital areas in most of the conditions. When comparing each recording condition to the beginning of the isolation period (Mar/ Apr), we found no significant differences in the distribution of theta power during the constant darkness period (May/Jun and Jul/Aug), nor at the end of the isolation period (Sept/Oct) (linear mixed-effects model with the isolation conditions as a fixed effect and different random intercepts for each participant, t-values, n=14 recordings at Noon and n=18 recordings in the Evening) (Fig. 3.b).

**Figure 3:**
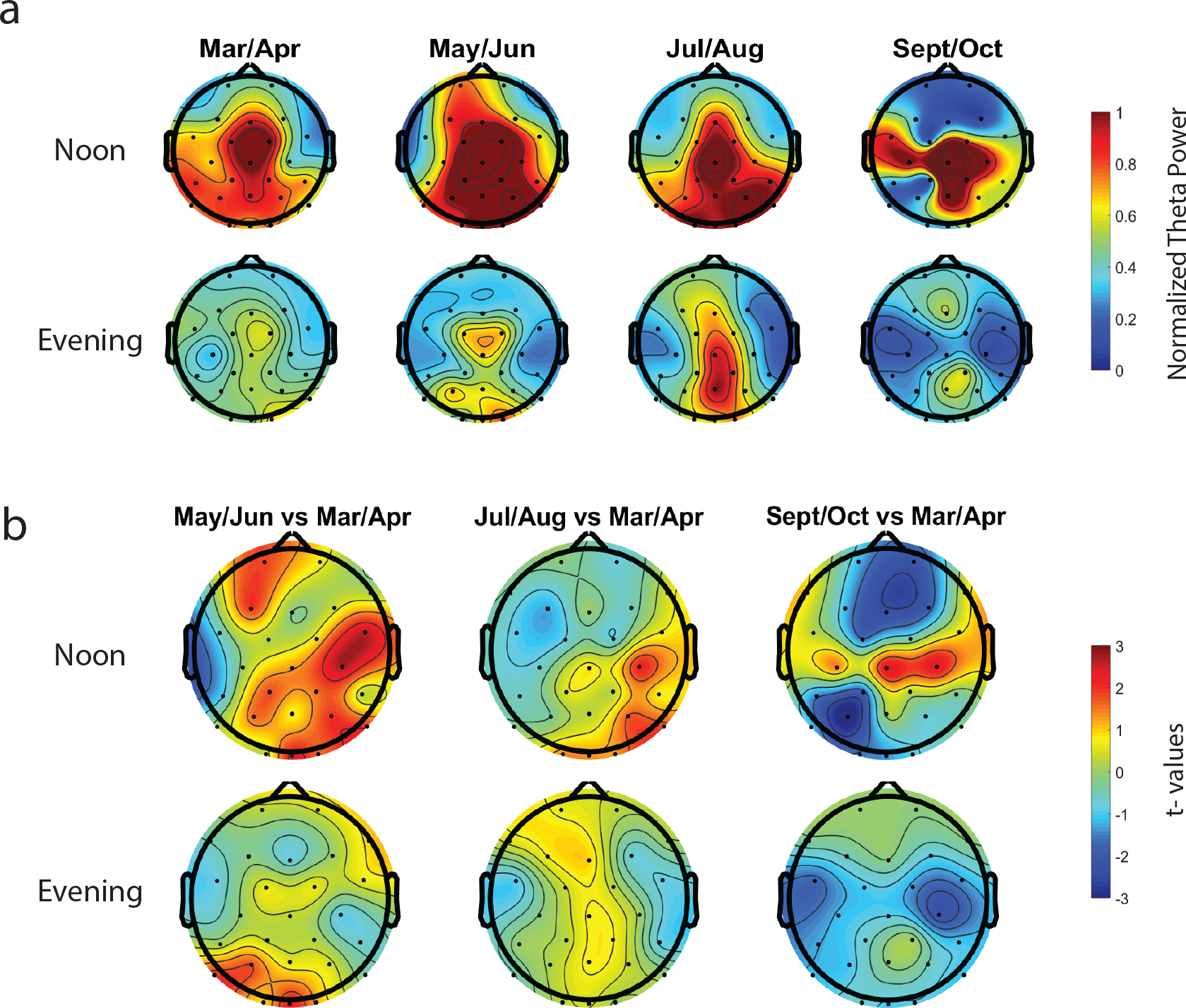
Theta power topographical distribution during the isolation period. We have 4 recordings per participant during the isolation period (Mar/Apr, May/Jun, Jul/Aug, Sept/Oct). (a) Normalised theta power per condition for recordings taken at Noon and in the Evening. (b) No significant differences in theta topographical distribution when comparing May/Jun, Jul/Aug and Sept/Oct to Mar/Apr

### Theta power, sleepiness and psychological strain

After each resting state EEG recording, the participants had to report their subjective psychological strain by rating adjectives on a questionnaire. We analysed the psychological strain scores’ relation with theta power at Noon and in the Evening. We first represented the relation between sleepiness and theta power for all electrodes over the scalp. We found that in the Evening sleepiness was high when theta power across the scalp was high. This was not the case in the Noon recordings. (linear regression model, t-values, white dots for uncorrected p-values<0.05, for a cluster to be significant it should contain at least two neighbouring electrodes, n=14 recordings at Noon and n=18 recordings in the Evening) (Fig. 4.a). Then, while looking at the psychological strain questionnaire’s total score, we found that only the right frontal cortex (electrodes F4 and F8), has high theta power when psychological strain scores are high (linear regression model, t-values, white dots for uncorrected p-values<0.05, for a cluster to be significant it should contain at least two neighbouring electrodes, n=14 recordings at Noon and n=18 recordings in the Evening) (Fig. 4.b). For theta power within the right frontal cluster (electrodes F4 and F8), we computed the best linear fit for its relation with the psychological strain scores (linear regression model, r^2^ = 0.363, p=0.008, n= 18 recordings) (Fig. 4.b). This correlation was not driven by any isolation condition in particular and seems to be a general trend.

**Figure 4:**
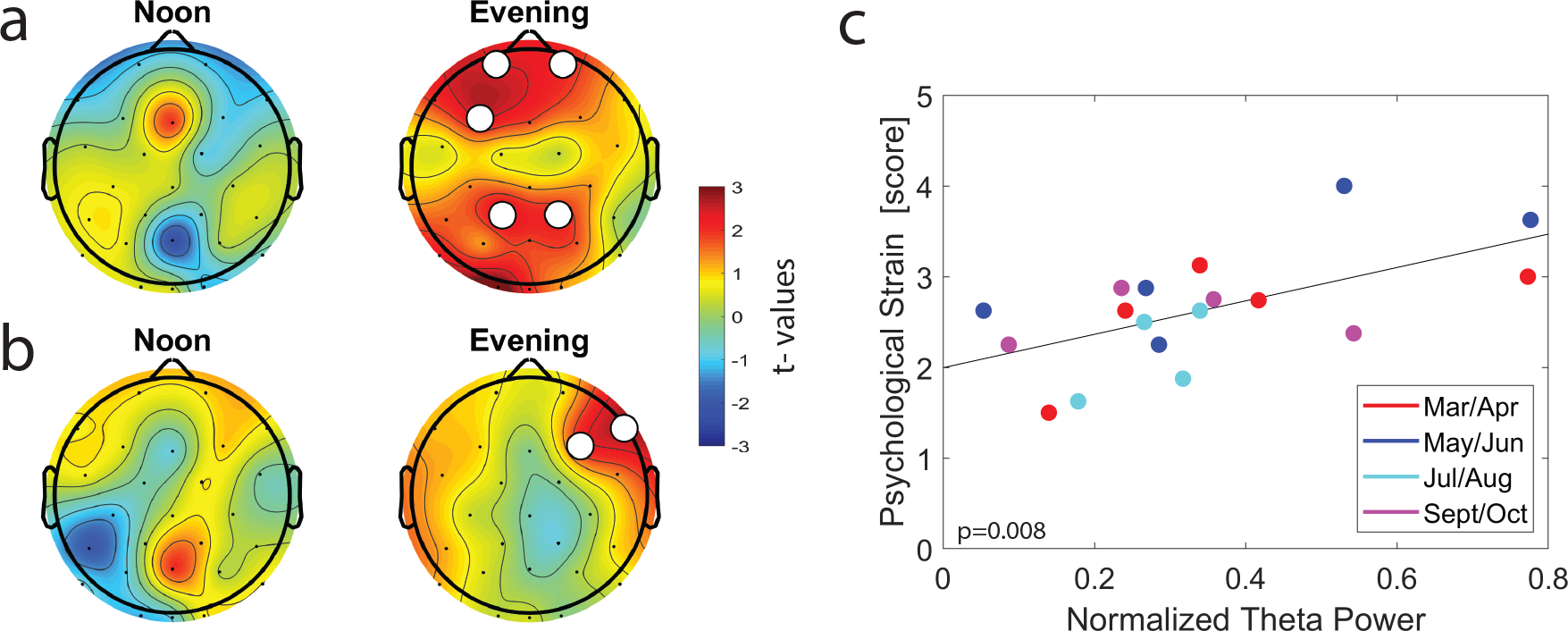
Relation between sleepiness, psychological strain and theta power. (a) Topographical representation of sleepiness as a predictor for theta power at Noon and in the Evening. Electrodes in the frontal central and parietal areas (FP1, FP2, F3, CP1, CP2, P7, O1 and O2) have a significant positive slope (white dots for uncorrected p-values<0.05). Meaning that when theta power in high sleepiness scores are high. (b) Topographical representation of subjective psychological strain as a predictor for theta power at Noon and in the Evening. Electrodes in the right frontal area (F4 and F8) have a significant positive slope (white dots for uncorrected p-values<0.05). Meaning that when theta power is high psychological strain score are low. (c) Positive slope for the relation between the psychological strain score and the mean theta power at electrodes F4 and F8 for all Evening recordings.

### Theta power and physical activity

Then, for approximately one hour (1.3*h* ± 0.1), each participant had to perform an incremental bicycle exercise. We compared the power spectrum at the electrode C3 Before and After the physical exercise and we found a reduction After the physical exercise in the Noon session in the 3.5-8.5Hz range (paired t-test, blue dots for uncorrected p-values<0.05, df=13, n= 14 Noon recordings, n=18 Evening recordings) (Fig. 5.a). When only comparing theta power (5-7Hz), we found a reduction After the physical exercise in the Noon session (paired t-test, t=−3.487, p=0.004, df=13, n=14 recordings). No such reduction was found in the Evening session (paired t-test, t=0.369, p=0.717, df=17, n=18 recordings) (Fig. 5.b). We applied the same approach for each electrode over the scalp and found that theta power was significantly reduced in the central, parietal and temporal areas After the physical exercise in the Noon session (paired t-test, t-values, white dots for uncorrected p-values<0.05, for a cluster to be significant it should contain at least two neighbouring electrodes, n=14 recordings at Noon and n=18 recordings in the Evening) (Fig. 6.b).

**Figure 5:**
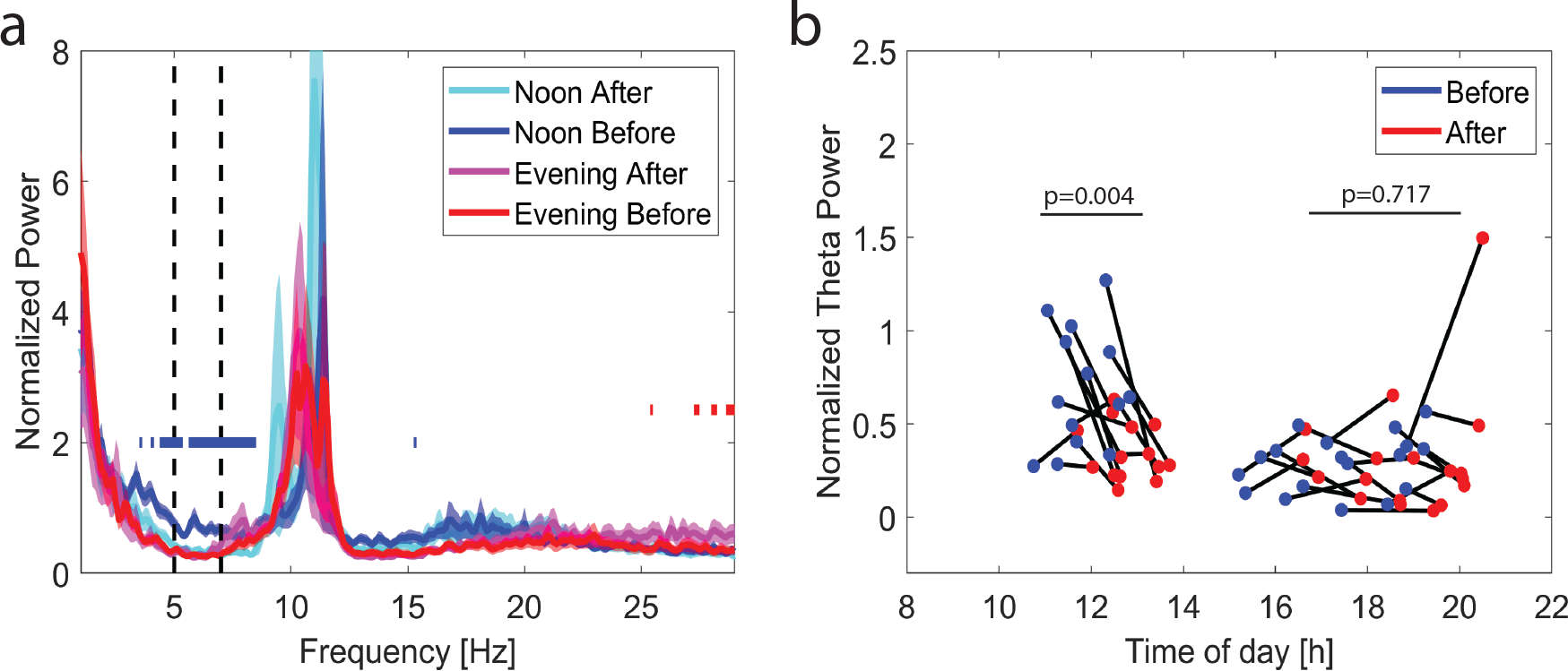
Physical exercise effects on theta power. (a) The power spectrum at electrode C3 Before and After the physical exercise for Noon and Evening recordings (*mean±sem*). Significant decrease of activity in the (3.5-8.5Hz) range After the physical exercise at Noon (blue dots for Noon and red dots for Evening’s uncorrected p-values<0.05). (b) The blue dots represent theta power at electrode C3 Before the physical exercise and the red dots represent theta power at electrode C3 After the physical exercise. There is a significant decrease of theta power at Noon after the physical exercise. No significant decrease of theta power in the Evening after the physical exercise.

**Figure 6:**
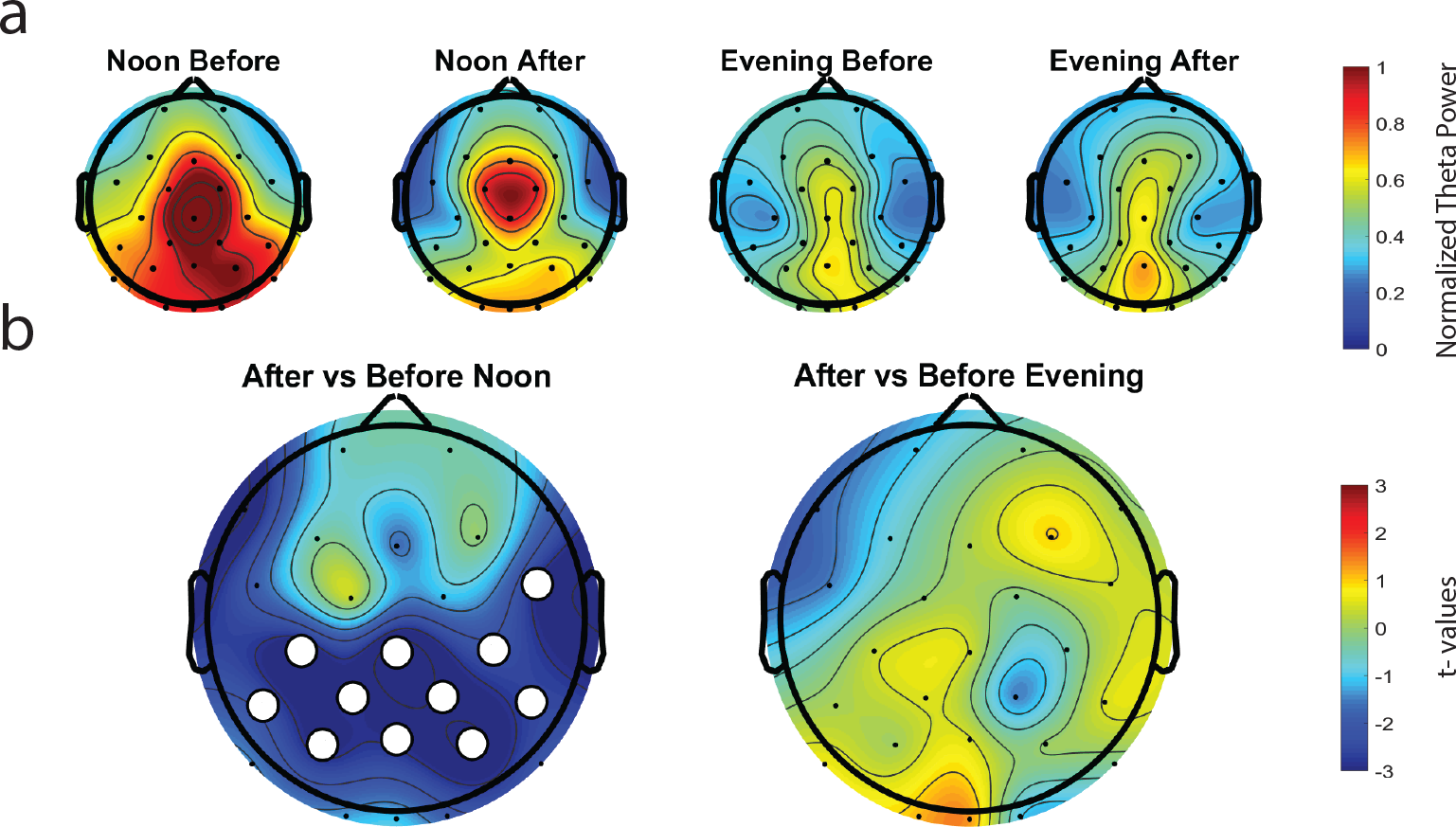
Theta power topographical distribution Before and After the physical exercise. (a) Normalised theta power at Noon Before and After the physical exercise and in the Evening Before and After the physical exercise. Consistent central, parietal and occipital distribution of the theta power in all conditions. (b) Decrease of theta power in the central, parietal and temporal areas after the physical exercise at Noon (white dots for the uncorrected p-values<0.05)

## Discussion

We found no differences in theta power across the isolation conditions. However, across conditions, we found that an increase of theta power in the evening in the right frontal area was associated with an increase of psychological strain. Finally, we found evidence that a bicycle exercise is able to reduce theta power, at least at its peak around noon.

Misalignment between the endogenous circadian clock and the sleep wake cycle reduce sleep quality and quantity (36) and chronic sleep restriction is known to increase sleep pressure (29). In a previous study, we showed an increase of sleep pressure markers in ISS astronauts (15). One possible explanation for such increased sleep pressure on the ISS might be long-term isolation. However, long-term isolation on Earth at the Concordia station, did not show any influence on electrophysiological measures of sleep pressure. Hence, isolation might not be the cause for increased sleep pressure on ISS. Eight months of isolation at the Concordia station was not sufficient to mimic the six months’ space mission on the ISS.

A review by Pagel and Chouker outlined that isolation can impact cognitive performances, stress levels and psychological strain (37). Our questionnaire was designed to assess psychological strain during the isolation period at the Concordia station. In our study, we showed an increase of subjective sleepiness when theta activity was high in the evening, which confirmed the role of theta power as a sleep pressure marker during wakefulness. Moreover, theta power had no correlation with sleepiness at noon, which confirmed the strong circadian regulation of sleepiness (38). In the evening, an increase of psychological strain was correlated with an increase of theta power in the right frontal cortex. Usually, theta power is low in the evening, since the circadian component has a strong influence on sleep pressure at this time of day (9; 11; 12). If the circadian rhythm is misaligned (i.e. phase shift), evening sleep pressure will not properly be compensated for. In turn, theta power would be higher than normal, which could impact sleepiness and psychological strain. The relation between circadian misalignment and mood is well known (39). However, in our study the correlation between theta power and psychological strain needs to be further explored in a laboratory controlled environment.

Overall, countermeasures need to be implemented to improve the sleep quality and quantity in further interplanetary space exploration missions. Daily physical exercise has shown to help maintain the psychological strain of the participants during the isolation period at Concordia station (32). Furthermore, Pattyn and colleagues outlined that daily physical exercise can help reduce sleep disturbances in Antarctica (23). From the oscillation of theta power during the day, we can expect a peak of theta activity at 14h (9; 11; 12). Accordingly, for recordings around noon, we would expect an increase of theta power after one hour of physical activity. However, in our study, we noticed a decrease of theta power after the bicycle exercise when the exercise was performed at noon. Our study is a first indication that an ergometer bicycle exercise could decrease sleep pressure in some cortical areas. At noon, the circadian component doesn’t compensate for the build up of sleep pressure. It is hence not surprising that only at this time of day we can manipulate sleep pressure with countermeasures. In the evening when the circadian component had a stronger influence on sleep pressure, physical exercise had no effect on sleep pressure markers. To confirm the interest of physical activity as a countermeasure for sleep pressure, further studies would need to record sleep EEG the following night to observe the SWA levels at the beginning of the night and study how SWA dissipates overnight.

In the work by Fisher and colleagues (35) it is hypothesised that wheel running is an automatic movement for mice, also called stereotypic movement. This type of exercise would allow a decrease of firing activity in some cortical areas. In our study, we hypothesise that bicycling in humans could have a similar effect and that an appropriate physical exercise could locally decrease sleep pressure. Even though, physical activity involving novelty and more complex tasks often show an increase sleep pressure (40). Previous research on the effects of physical activity on cortical activity, revealed that cortical activity was specific to the type of physical exercise, that there is a familiarisation process with the exercise and that there is an intensity response to physical activity (41). For example, at high intensity and when the subject are familiar with the exercise, a reduction of beta activity was observed in the prefrontal cortex during resting state with closed eyes (42). During the MARS 500 space analogue mission, cortical activity in the frontal areas was decreased and cortical activity in the parietal lobe was increased after physical activity involving endurance (i.e. two 30 min session of treadmill running and ergometer bicycling) in resting state EEG with closed eyes (43).

Laboratory condition can unfortunately not simulate properly spaceflight conditions. Concordia station shares several psychological and physical aspects of human space exploration and is one of the most remote space analogue facility. Nonetheless, to understanding the mechanisms underlying sleep disturbances in Antarctica and test further sleep pressure countermeasures, we would need a more controlled environment. The station medical doctors, reported night shifts during the winterover for some participants. The main limitation of this study is the lack of information on the sleep-wake history during the isolation period. The amount of sleep the night before the recordings and the wake up time were not reported. Accordingly, we cannot assume that all participants are aligned with their circadian clock. In further experiments, a better controlled sleep-wake schedule should be implemented to study isolation’s effect on sleep pressure. In addition to wake EEG data, sleep EEG data and sleep-wake history would have been necessary to confirm our findings. In a previous study, an increase of beta activity in the prefrontal cortex was observed after an acute hypoxic exposure (44). Even though a fast adaptation was shown (16), we cannot exclude that hypoxia might affect our results. In this study we had no baseline recordings before participants’ trip to Antarctica. As a consequence, we used the recording at the beginning of the isolation period (Mar/Apr) as a baseline to study the other isolation recordings. In Mar/ Apr the temperatures are not at their minimum, the summer camp crew just left the station and the winter crew is preparing for the winterover. Similarly, the Sept/Oct isolation recordings represents the end of the isolation period, since the sun rise again and planes start coming back to the station. For each participant and along the winterover, experiments were performed at different times during the day. To compare recordings within participants, we had to exclude about 1/4 of the recording sessions when recorded at ±1*hour* from the other recordings. Because of this exclusion criteria we used a mixed effects model to compensate for missing recordings. In EEG studies, it is common to observe inter-individual differences between participants (45). Hence, it is not ideal to compare

Noon recordings with Evening recordings with different participants in each group. Moreover, participants were recorded over two years (DC7 and DC8), which could increase inter-individual differences.

Taken together, our results suggest that increased sleep pressure in the evening impacts psychological strain and physical activity at noon could be envisioned as a countermeasure for high sleep pressure.

## Methods

### Participants and experiment

12 participants (all male, 40 ± 0 years old) took part in the experiment (ESA Experiment record n° 9363) over the winter 2011 and 2012 (winterovers DC07 and DC08) at the French-Italian Concordia research station in Antarctica (Fig. 1.b). Written informed consents were obtained prior to participation. The experimental protocol was approved by the European Space Agency’s Medical Board (ESA-MB) and the ethic committee from the Institut Polaire Paul Emile Victor (IPEV). During the period of isolation, the participants took part in the experiment once every six weeks. During each recording session, the participants had a five minutes resting state EEG recording with eyes open and they had to answer to a psychological strain questionnaire. This was followed by an incremental bicycle exercise lasting 1.3 ± 0.1 hour for each session. Finally, a second five minutes resting state EEG, with eyes open, was recorded. Also 5 minutes of wake resting EEG with eyes closed was recorded just after each open eyes recordings. We decided to only analyse the open eyes data since, theta power during closed eyes doesn’t correlate with sleepiness (46). In this study, we defined the recording conditions for each session during the isolation period with the labels Mar/Apr, May/Jun, Jul/Aug and Sept/Oct. A training session to get familiar with the tasks was performed at the Concordia station three weeks before the beginning of the winter period (i.e. winter between Mar/Oct). However, these recordings were not used in this analysis. Out of the 48 recording sessions, four sessions were not well recorded and excluded from the dataset. Recordings took place at variable times along the day. We excluded all sessions performed at more than ±1 hour from the median time of recording throughout Mar/Apr, May/Jun, Jul/Aug and Sept/Oct sessions for each participant. Out of 44 sessions, 12 sessions were excluded from further analysis because of this lack of circadian similarity between the isolation sessions. During the winter’s constant darkness period (i.e. May/Aug), the temperature drops rapidly toward a mean temperature of −60° with peaks at −80°C (Fig. 1.a).

### Wake EEG recordings

Each participant, for each session, was recorded with a 32 channels actiCAP (BrainProducts, Gilching, Germany) (10–20 electrode system EEG cap). Continuous wake EEG with eyes open was recorded for 5 minutes during each session at a sampling rate of 500 Hz and down sampled to 250Hz. Scalp electrodes’ impedance were measured and kept below 10 KΩ. All channels were referenced to FCz. ECG and electrodes TP9 PO9 Po10 T7 and T8 were excluded from further analysis to standardise DC07 and DC08 recordings. Out of the 32 electrodes recorded, 26 remained in our analysis. EEG data pre-processing was performed in Matlab (Version R2018b) using EEGLAB toolbox scripts (Version 14) (47) and additional custom made scripts. EEG data were down sampled to 250 Hz and pass-band filtered [0.1–48 Hz]. Each EEG channel was referenced to the average activity across the channels (i.e. average referencing). An Independent Component Analysis (48) was performed to remove ocular, muscular, and electrocardiographic artifacts (3.12 ± 0.18 components rejected per recording) as defined by Hulse and colleagues (49). Using the EEGLAB graphical user interface, all movement’s artefacts in the signal were marked by visual inspection and removed. The power spectrum was computed for each channel and outliers containing high muscle artefacts (20-30Hz) were excluded from the dataset (1.89 ± 0.15 channels rejected per recording) (50). For each subject, rejected channels were interpolated. To reduce the signal to noise ratio, we performed a phase-rectified signal averaging (PRSA) (51). PRSA allows to superimpose the oscillations to create an interference and hence reduce the weight of acute noise generators in the signal. The power spectral density was estimated using the Welch’s averaged periodograms with a four second Hamming window and a frequency resolution of 0.125 Hz. In each frequency bin, the power at each channel was normalised by the average power over the scalp. The theta power band was computed between 5 and 7 Hz.

### Psychological strain questionnaire

The participants completed a questionnaire measuring their psychological strain. This questionnaire is derived from the “Eigenzustandsskala” (EZ-K, which is translated as “Personal state scale”) (52). The questionnaire measured the subjective psychological strain by asking the participants to rate their sleepiness, psychological strain, calmness and recovery state between zero and five. In our study, we reported the psychological strain questionnaire score, rated between zero and five. Since the questionnaire had one question about sleepiness we extracted this unique variable and reported the score, also rated between zero and five. Five being the highest level of sleepiness and zero being the lowest. The questionnaire was presented in the participants’ native language.

### Physical exercise

The particpant had to perform an incremental bicycle ergometer exercise. The physical exercise started with a 50 Watts workload and increased by 30 Watts every 3 minutes until subjective exhaustion of the participant (lasting 1.3 ± 0.1 hour for each session).

### Statistics

In all statistical analysis we first assessed normal distribution of the data with a quantile-quantile plot. When comparing measures in different participants, for example when comparing the Noon and the Evening sessions, we performed a two sample t-test (t=t-value, p=p-value, df=degrees of freedom). When comparing measures within participants in two different conditions, for example when comparing theta power Before and After the physical exercise, we performed a paired t-test (t=t-values, p=p-value, df=degrees of freedom). When studying repeated measures within the same participants, for example when comparing isolation conditions, we performed a linear mixed effects analysis for the relationship between the response variable and the fixed/random effects. We used the restricted maximum likelihood estimate method to fit the model and choose the best model based on Bayesian information criterion results. Visual inspection by quantile-quantile plot of the residuals confirmed that homoscedasticity and normality were respected. Then, we completed a two-sided t-test (t=t-value, p=p-value, df=degrees of freedom) with the null hypothesis that each fixed effect in the model has a coefficient of zero. When the best model to study the response variable contain only one fixed effect without random effects, for example when looking at the relation between theta power and psychological strain scores, we used a linear regression model to obtain a R^2^ value which indicates how much of the total variation can be explained by the fixed effect. Then, we completed a two-sided t-test (t=t-value, p=p-value, df=degrees of freedom) with the null hypothesis that the slope of the model is equal to zero. For the topographical representations we plotted the t-values for the aforementioned t-tests and we draw white dots for significant p-values (uncorrected p-values<0.05). Then, we completed a non-parametric permutation test (critical value=2.14 at Noon or 2.10 in the Evening, 500 permutations, n=14 recordings at Noon session and n=18 in the Evening) for multiple testing correction (40; 53), defining the minimum cluster size for a pattern to be significant. In our analysis at least 2 neighbouring electrodes needed to be significant to be reported as a significant effect. All statistical analysis were performed in Matlab.

### Data availability

All relevant data will be available from the corresponding authors upon request and after approval from the European space Agency Medical Board (ESA-MB).

### Code availability

All relevant scripts will be available from the corresponding authors upon request.

## Acknowledgements

This work was supported by the research grant 50WB0819 from the German Space Agency (DLR) and by a research grant from the University of Zurich (Clinical Research Priority Program Sleep and Health). It was also supported by the European Space Agency (ESA) and the Luxembourg Government.

## Author contributions

Abeln and S. Schneider conceived the experiment. G. Petit, R. Huber and L. Summerer designed the data analysis study. G. Petit performed the data analysis and wrote the manuscript. R. Huber gave data analysis support. All authors contributed to the scientific discussion and manuscript revisions.

